# Optimisation of asymmetric field pulses for transcranial magnetic stimulation

**DOI:** 10.1101/2023.05.01.538933

**Authors:** Ke Ma, Stephan M. Goetz

## Abstract

Transcranial magnetic stimulation (TMS) is a widely-used noninvasive brain stimulation technique through electromagnetic induction. Nowadays commercial TMS devices routinely use conventional biphasic pulses for repetitive TMS protocols and monophasic pulses for single-pulse stimulation. They respectively generate underdamped or damped cosinusoidal electric field pulses that have been proven to be power-inefficient. Recently, symmetric field pulses with near-rectangular electric fields show great potential in terms of energy loss and coil heating, but only limited studies have investigated asymmetric field pulses with different asymmetry levels for the induced electric field waveforms. Thus, this paper optimises and searches a wide range of potential current waveforms with the goal of minimising energy loss and coil heating. The optimised results demonstrated that asymmetric field pulses with near-rectangular electric fields have significantly lower power consumption than conventional ones. These optimised waveforms commonly consist of an initial falling phase followed by rapidly rising and falling phases, trailed by a slow decay to zero. Interestingly, the initial phase has a decay time constant around 260 µs and introduces a pulse-duration-dependent negative bias for the current baseline to minimise the energy loss and coil heating. Thus, it is possible to directly design asymmetric field pulses with various asymmetry ratios by using several prediction equations rather than running optimisation. These results also suggest that introducing such an initial current phase could likely significantly reduce the coil heating of most TMS pulse shapes to improve their power efficiencies.

## 1 Introduction

Transcranial magnetic stimulation (TMS) is a widely-used noninvasive brain stimulation technique through electromagnetic induction (Barker et al. 1985; Polson 1982). Through large alternating currents in a stimulation coil, TMS generates short magnetic field pulses that can easily penetrate the skull and evoke action potentials of cerebral neurons in the focus of the specific coil (Hallett 2000). During a pulse, typically no longer than a millisecond, the stimulation coil produces magnetic flux densities in the low Tesla range and induces electric fields (E-field) of some 100 V*/*m. The currents reach thousands of amperes, requiring substantial power and causing rapid heating of the device and the stimulation coil (Goetz and Deng 2017; Wagner et al. 2007). The high power requirement is widely responsible for the inflexibility of available devices, complicates device technology, requires active cooling systems for more advanced stimulation protocols, and still limits TMS sessions (Nielsen et al. 1995; Rossi et al. 2021, 2009)

Depending on the direction of the induced E-field inside the brain tissue, TMS pulses can be classified into monophasic (i.e., asymmetric and more unidirectional E-field) and biphasic (i.e., more symmetric and thus more bidirectional E-field) families, which are widely available in commercial devices and routinely used (Kammer et al. 2001; Rossi et al. 2021; Rossini et al. 2015). Conventional monophasic stimulators generate an initially cosinusoidal E-field pulse that terminates through massive overdamping with a resistor, which turns the entire pulse energy into heat (Polson 1982), whereas biphasic pulses have almost cosinusoidal E-field pulses that typically end after one period, half a period (half-sine) or, more rarely, multiple periods (Cadwell 1991; Peterchev et al. 2015a). Without a damping resistor for pulse shaping, the oscillator circuit can recover a large portion of energy, typically even the majority of the pulse energy after each pulse. Therefore, biphasic pulses can be generated more efficiently and are almost exclusively used in high-frequency repetitive TMS, while monophasic pulses serve primarily in single-pulse and low-frequency stimulation (Goetz and Deng 2017; Rossini et al. 2015).

Monophasic pulses have advantages in stimulation selectivity and neuromodulatory efficacy (Arai et al. 2007; Goetz et al. 2016; Kammer et al. 2001; Niehaus et al. 2000; Sommer et al. 2006, 2002; Taylor and Loo 2007). For instance, Di Lazzaro et al. (1998, 2001b) reported that the recruitment of descending volleys is highly sensitive to the direction of the induced E-field for monophasic pulses. In contrast, biphasic pulses demonstrate less directional effects on the cortical output (Di Lazzaro et al. 2001a, 2012). In consequence, monophasic pulses lead to different response latencies in response to stimulation in the primary motor cortex, which can furthermore be controlled through the direction of the induced E-field. These results indicate that monophasic pulses can selectively activate a relatively uniform population of neurons. Furthermore, several studies have clearly demonstrated that monophasic repetitive TMS is more effective than biphasic ones (Arai et al. 2007; Sommer et al. 2002; Taylor and Loo 2007; Tiksnadi et al. 2020). The latency varying from 1 to 1.5 ms is assumed to be on the order of a synaptic transmission delay and is often associated with stimulating more upstream neural populations (Sommer et al. 2006).

While conventional regular-rate repetitive TMS tends to be more effective with monophasic pulses at least in the primary motor cortex, some paradigms such as quadri-pulse stimulation require and seem to primarily work due to monophasic pulses (Hamada et al. 2007). Quadri-pulse stimulation with biphasic pulses has a much shorter after-effect duration than that with monophasic pulses (Nakamura et al. 2016). Due to the apparently higher efficacy, monophasic pulses are profoundly attractive for clinical therapy. However, at present monophasic devices are highly inefficient and cannot provide higher rates. Furthermore, the coil heating of such pulses should be as low as possible for repetitive protocols. To resolve this problem, researchers have explored various solutions, including active cooling systems and coil optimisation (Koponen et al. 2015, 2017; Lin et al. 2000; Nielsen et al. 1995; Vachenauer 1999; Wang et al. 2018). However, the technical effort and the usability disadvantages, particularly of cooling, can be high, and most of the potential of coil design has already been exploited.

If not generated through conventional technology, monophasic pulses are slightly less efficient than cosinusoidal biphasic ones, but far behind rectangular pulses (Goetz et al. 2012a, 2013, 2012b; Niehaus et al. 2000). Biphasic pulses with symmetric E-field shape, however, were optimised through numerical optimisation with nonlinear neuron models to develop pulse shape features with higher efficiency (Goetz et al. 2013). Most recent research suggested a more efficient monophasic pulse by only optimising the waveform of the original monophasic electric field pulse without considering the overall pulse waveform (Wang et al. 2023). The full optimum of asymmetric field pulses, ideally for different asymmetry ratios is missing.

To address this research gap, we aimed to optimise the parameters of the current waveforms of asymmetric field pulses to reduce energy loss and coil heating. Thus, we established an objective function of energy loss during magnetic stimulation for the asymmetric field pulses and used a user-defined optimisation algorithm to adjust the current waveform without constraining it. Finally, our study demonstrates the major advantages of the optimised asymmetric field pulses over standard monophasic pulses.

## 2 Methods

### Objectives and Constraints

We minimise the energy loss during a pulse, which determines the required supply power and heats the coil. The amount of heat generated in the coil is directly proportional to the energy loss, which, in turn, limits the maximum duration of the stimulation session and the pulse repetition rate (Riehl et al. 2008; Rossi et al. 2009; Weyh et al. 2005). The losses

that heat up the device and the coil are dominated by Joule heating and can be described through the internal resistance in the first approximation. For a coil current *i*(*t*), the energy loss of a pulse is

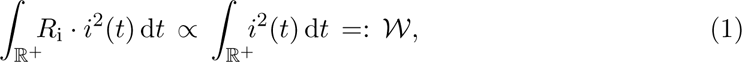

where *R*_i_ *>* 0 is the equivalent intrinsic resistance. As the internal resistance is normally independent of the pulse shape, the optimisation excludes this proportionality constant and uses the functional *W* in the following.

We impose two important constraints on the optimisation problem for asymmetric field pulses. First, the induced E-field pulse, which is proportional to the time derivative of the coil current 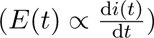, must be of sufficient strength to successfully evoke an action potential in a neuron. Second, we limit the maximum positive and minimum negative voltages, *V*_max_ and *V*_min_, forming a voltage and E-field asymmetry ratio *m* = *r*_E_ = *r*_V_ = 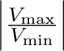. The E-field ratio of positive versus negative pulse contributions furthermore appears to cause the selectivity property of monophasic or more generally asymmetric field pulses (Halawa et al. 2019; Sommer et al. 2018). The voltage limit further reflects the practical limits of available devices as well as device concepts due to ratings of electronic components and safety considerations; furthermore, they are also justified by the test requirements in medical device regulation (Goetz and Deng 2017; Rossi et al. 2021). Without voltage limits, the pulse optimisation will shorten the pulse at the cost of higher voltages as well as peak electric fields, which are known to reduce the energy loss even without changing the pulse shape, and accordingly diverge to extreme voltage values (Barker et al. 1991; Goetz et al. 2012b; Peterchev et al. 2013; Weyh et al. 2005). However, shorter pulses in the form of scaling a known pulse have limited scientific experiments while it further changes properties such as scalp sensation, the pulse sound, and apparently also the preferentially activated neural elements (D’Ostilio et al. 2016; Goetz et al. 2014; Peterchev et al. 2017, 2015b).

The optimisation problem identifies a function *i*(*t*): ℝ *→* ℝ representing the coil current over time such that the energy loss is minimal while ensuring the induced E-field pulse can elicit an action potential in neurons. The current function, *i*(*t*), is parameterised to translate the variational problem to a numerical optimisation procedure. The parametrisation of the current shape uses splines, i.e., local differentiable low-order functions, whose degrees of freedom are handed over to an optimisation algorithm. The degrees of freedom are mathematically equivalent to the number of anchor points on the curve. This representation furthermore allows the calculation of the voltage and the electric field time courses, which are linked to the current through the derivative. This parametrisation translates a set of parameters with finite but variable degrees of freedom *n* into continuous current profiles. It is stable and general so that any waveform can be approximated by a sufficiently high value of *n*. Furthermore, the degree of freedom *n* can be varied dynamically to refine solutions and allow for higher-frequency features, and existing solutions with one number of degrees of freedom can be converted to another one.

Furthermore, we translated the constraint of the positive and negative voltage limits into a penalty rather than fragmenting the solution space to allow for a potentially faster return to the valid solution space by presenting a steep but well-defined gradient or slope back once the optimisation exceeds the voltage limits. Each valid pulse further needs to cause a supra-threshold neuronal activation of a human nonlinear neuron model as used in previous TMS work in the form of an action potential (Cazenave-Loustalet et al. 2007; Goetz et al. 2013; Motz and Rattay 1986). The stimulation coil voltage is proportional to the time derivative of the coil current 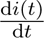. The objective function can accordingly be written as

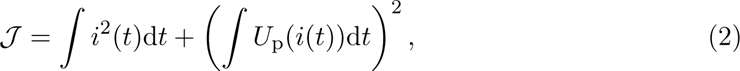

where *U*_p_ is the penalty function for the voltage when it exceeds the lower or higher voltage limits, which follows

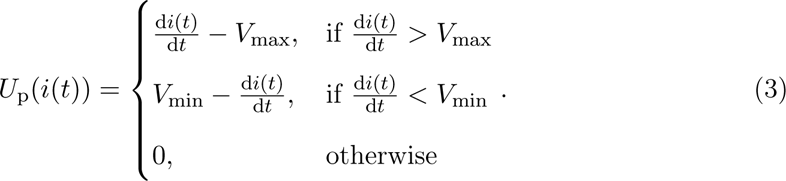

### Optimisation Procedure

As the current function is finitely described piece-wise by local curves, their parameters act as degrees of freedom for the optimisation algorithm. The optimisation approach uses a hybrid global–local framework since the problem is highly nonlinear, and there are numerous local minima. The local optimisation algorithm uses an interior-point method, which can rapidly converge to dominant local minima (Byrd et al. 2000). The local optimiser instances are nested inside a global particle-swarm optimiser, which in each iteration starts several parallel local optimisers. The method adaptively adjusts the number of degrees of freedom based on the improvement of the solution in the previous iteration step of the global optimiser as previously established (Goetz et al. 2013).

We implemented the particle swarm method to combine the information about the local minima found by the local optimiser instances and update the waveform parameters for them (Eberhart 1995). We used *ω* = 0.9, *c*_1_ = 1.2, and *c*_2_ = 0.12 for the particle swarm method, where *ω* is the inertia, *c*_1_ and *c*_2_ are the gravity parameters. The number of degrees of freedom is controlled by the global framework. The degrees of freedom are increased by a predefined factor (i.e., 5 per iteration) if convergence is achieved in the current iteration and the result outperforms the previous one; otherwise, the number of degrees of freedom is decreased. A new global minimum is updated if a local minimum has a lower cost compared to the current global minimum. The method was set up in Matlab (R2022a, Mathworks, USA). The waveform time window for the optimisation was set to 3 ms, and the time step for the spline was set to 1 µs.

### Data Analysis

We tested different combinations of voltage limits and asymmetry ratios, ranging from *m* = 0.25 to *m* = 20, with the highest limit being *V*_max_ = 2000 and the lowest *V*_min_ = *−*2000. The number of degrees of freedom reached up to one thousand during the optimisation process. The induced E-field pulses or coil voltage pulses with *m ≈* 1 are called symmetric field pulses in the following. To ensure a consistent comparison between the symmetric and asymmetric field pulses, we conducted optimisation for both.

The important parameters are all listed in Table 1. As shown in Figure 1, a typical current waveform has four distinct phases: a long initial phase, a steep rising phase, a similarly steep falling phase, and a shallow decay phase. Voltage limits show the upper and lower limits used in the penalty term in the objective function 2. *V̂*_max_ and *V̂*_min_ are respectively the true maximum and minimum coil voltages of the optimised results. *τ*_init_ is the time constant fitted by an exponential function, 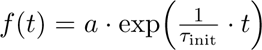, for the initial phase of the current waveforms. For simplification, we defined the effective pulse duration, *T*_pulse_, as the sum of the second (rising phase, *T*_rise_) and third (falling phase, *T*_fall_) current phases, where the coil voltage reaches at least 90 % of the true maximum *V̂*_max_ or minimum *V̂*_min_ coil voltages. *Î*_max_ and *Î*_min_ are respectively the maximum and minimum amplitudes of the optimised current waveforms. Moreover, we introduce the current ratio as 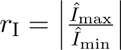 to evaluate the optimised current waveforms.

**Table 1:** The parameters of the optimised current waveforms and the corresponding voltage waveforms.

| Voltage limits | Voltage Parameters |  | Current Parameters |  |  |  |  | Energy loss and coil heating (a.u.) |
| --- | --- | --- | --- | --- | --- | --- | --- | --- |
| | $\hat{V}_{\max}$ (a.u.) | $\hat{V}_{\min}$ (a.u.) | $\tau_{\text{ini}}$ ($\mu\text{s}$) | $T_{\text{rise}}$ ($\mu\text{s}$) | $T_{\text{fall}}$ ($\mu\text{s}$) | $\hat{I}_{\max}$ (a.u.) | $\hat{I}_{\min}$ (a.u.) | |
| V:-100/+1000 | 995.0 | -99.9 | 254.0 | 77.0 | 503.0 | 42.3 | -34.9 | 469.8 |
| V:-100/+1500 | 1492.4 | -100.1 | 256.0 | 47.0 | 504.0 | 42.5 | -33.1 | 453.3 |
| V:-100/+2000 | 1990.0 | -99.8 | 264.0 | 36.0 | 409.0 | 43.4 | -30.7 | 446.4 |
| V:-250/+1000 | 995.0 | -248.9 | 260.0 | 81.0 | 212.0 | 54.5 | -26.8 | 379.2 |
| V:-250/+1500 | 1492.5 | -248.9 | 252.0 | 52.0 | 210.0 | 54.9 | -23.8 | 366.3 |
| V:-250/+2000 | 1990.1 | -248.8 | 264.0 | 37.0 | 212.0 | 54.8 | -24.9 | 357.5 |
| V:-500/+1000 | 995.0 | -497.6 | 258.0 | 85.0 | 119.0 | 63.7 | -22.5 | 346.6 |
| V:-500/+1500 | 1492.5 | -497.6 | 263.0 | 55.0 | 125.0 | 65.3 | -21.3 | 325.3 |
| V:-500/+2000 | 1990.0 | -497.5 | 264.0 | 41.0 | 125.0 | 66.4 | -18.0 | 314.8 |
| V:-1000/+500 | 497.7 | -995.0 | 250.0 | 187.0 | 53.0 | 65.9 | -27.1 | 405.5 |
| V:-1000/+1000 | 995.1 | -995.0 | 262.0 | 92.0 | 72.0 | 72.1 | -20.7 | 326.5 |
| V:-1000/+2000 | 1990.1 | -995.0 | 265.0 | 46.0 | 69.0 | 77.0 | -16.7 | 286.7 |
| V:-2000/+500 | 497.6 | -1989.1 | 250.0 | 188.0 | 26.0 | 70.3 | -25.8 | 399.6 |
| V:-2000/+1000 | 995.1 | -1990.1 | 261.0 | 99.0 | 32.0 | 78.9 | -20.4 | 314.5 |
| V:-2000/+2000 | 1990.1 | -1990.1 | 274.0 | 51.0 | 38.0 | 86.7 | -16.0 | 268.2 |

**Note:** (a) Voltage limits show that voltage constraints for the optimisation. For example, V:-100/+1000 means that the minimum and maximum limits are respectively -100 and 1000; (b) Voltage parameters, $\hat{V}_{\max}$ and $\hat{V}_{\min}$ , are the true maximum and minimum values of the optimised results; (c) $\tau_{\text{ini}}$ is the decay time constant for the initial phases ( $R^2 \geq 0.99$ for all of them); (d) $T_{\text{rise}}$ and $T_{\text{fall}}$ are respectively the duration of the rising and falling phases of current waveforms; (e) $\hat{I}_{\max}$ and $\hat{I}_{\min}$ are respectively maximum and minimum values of the optimised current waveforms; (f) Energy loss is the value of the Equation 1.

**Figure 1:**
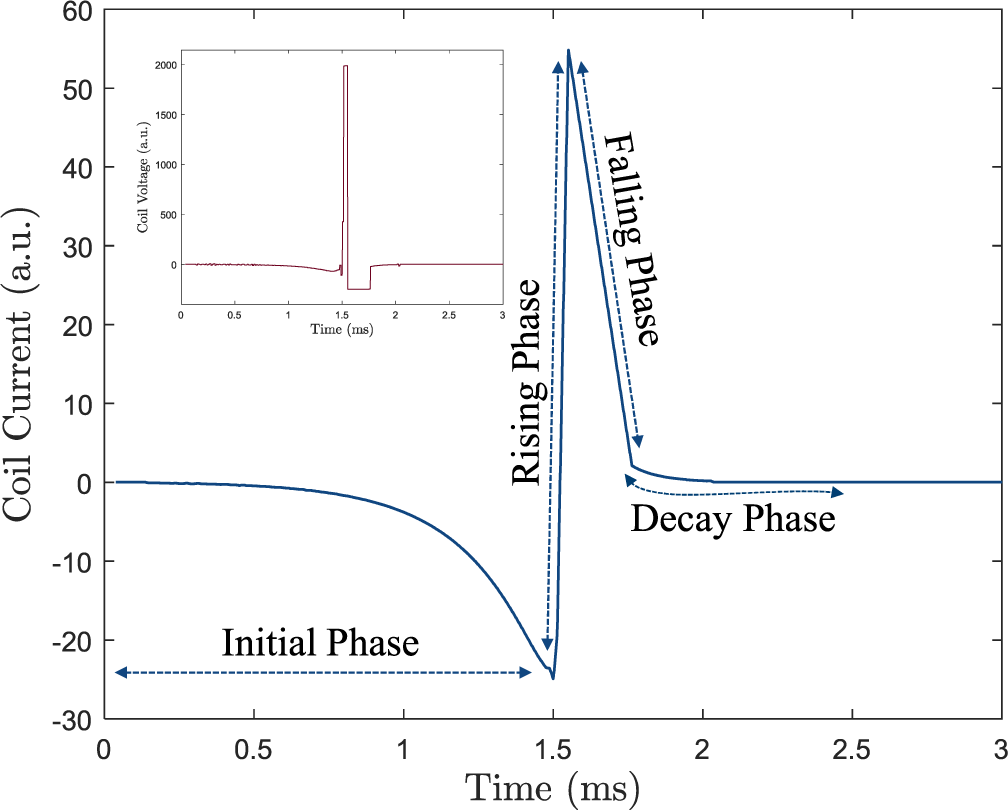
Coil current and voltage waveforms with *V*_max_ = 2000 and *V*_min_ = *−*250; a.u. denotes arbitrary unit. The current waveform has four distinct phases, specifically an initial phase, a rising phase, a falling phase, and a decay phase. The rising and falling phases form a triangular current and thus rectangular electric field waveform (see the inset). Voltage and electric field are determined by the inductive component of the coil and are directly proportional to the derivative of the current.

In the *Results* section, we showed the common features among the optimised cur-rent and voltage waveforms and investigated the relationships among the pulse duration, the current and voltage waveforms, and the energy loss calculated by the first item in Equa-tion 2. These relationships were fitted by a power function *f* (*X*) = *a · X^b^*, an exponential function *f* (*X*) = *a ·* exp(*b · X*), or a linear function *f* (*X*) = *a · X* + *b*. The corresponding coefficients of determination (*R*^2^) for each regression were also reported. Finally, we compared the optimised monophasic pulses given in Wang et al. (2023) with our results. All current waveforms are aligned in the following plots by shifting the minimum current value *Î*_min_ to the time point 1.5 ms.

## 3 Results

### The optimised waveforms have common features

The optimised current and voltage waveforms show similar patterns overall. Figure 1 displays one of the optimised monophasic current waveforms, and all of these current wave-forms typically have four distinct phases: a long initial phase, a steep rising phase, a similarly steep falling phase, and a shallow decay phase. The current amplitude slowly decreases to a negative value in the initial phase, then rapidly increases to the highest positive amplitude in the rising phase, followed by the falling phase, and then slowly decreases to zero in the decay phase. The corresponding coil voltage waveform is near-rectangular and asymmetric as shown in the inset plot in Figure 1. The unequal positive and negative voltage phases respectively correspond to the ascending and descending slopes for the rising and falling phases in current waveforms.

According to Equation 1, the energy loss is directly proportional to the squared current. The initial phase only slightly increases the energy loss as its current, and, particularly, the squared current is relatively low compared to the rising and falling phases. However, the initial phase can reduce the energy loss by introducing a negative bias for the subsequent rising phase, which pulls the peak current of the rising phase downwards. Thus, this negative bias decreases the energy loss by lowering the peak current in the rising phase without reducing the pulse duration. Interestingly, all optimised current waveforms have similar decay constants *τ*_init_ for the initial phases, ranging from 250 µs to 274 µs (*R*^2^ *≥* 0.99 for all fitted exponential functions).

Furthermore, it is worth noting that the effect of the initial phase on neural dynamics is minimal. According to the neural response, the membrane potential is slightly hyper-polarised at the end of the initial phase. The rising phase subsequently depolarises the membrane potential and brings it above the neural firing threshold of the sodium channels. Afterwards, the falling and decay phases hyper-polarise and counteract the activation. The last phase, i.e., the decay phase most likely delays the current decay and thus reduces the electric field to keep the threshold of the overall pulse low with marginal extra cost in loss or heating. Thus, the initial phase of the optimised pulses reduces the energy loss generated by the rising current phase and thus the primarily depolarising electric field phase, the third phase brings down the current as quickly as possible to more moderate currents with lower loss using the minimum available voltage (i.e., the lower voltage limit, *V*_min_), while the last phase can decay slowly for lower electric field strength and lower hyper-polarisation impact.

### Larger voltage amplitudes make voltage pulse shorter

The resulting voltage waveforms, shown in Figures 2 and 3, were almost rectangular with varying pulse widths. These plots suggest that a higher voltage amplitude would result in a shorter pulse for both the positive and negative voltage phases as expected (Peterchev et al. 2013). To investigate this relationship further, a scatter plot was created for *T*_rise_ versus *V̂*_rise_ and *T*_fall_ versus *V̂*_fall_ as shown in Figures 4 (A) and (B). These results indicate that the pulse widths for both positive and negative voltage phases decrease as the magnitudes of the pulses increase. The fitting curves describe these trends well with sufficiently high coefficients of determination (*R*^2^ *≥* 0.95).

**Figure 2:**
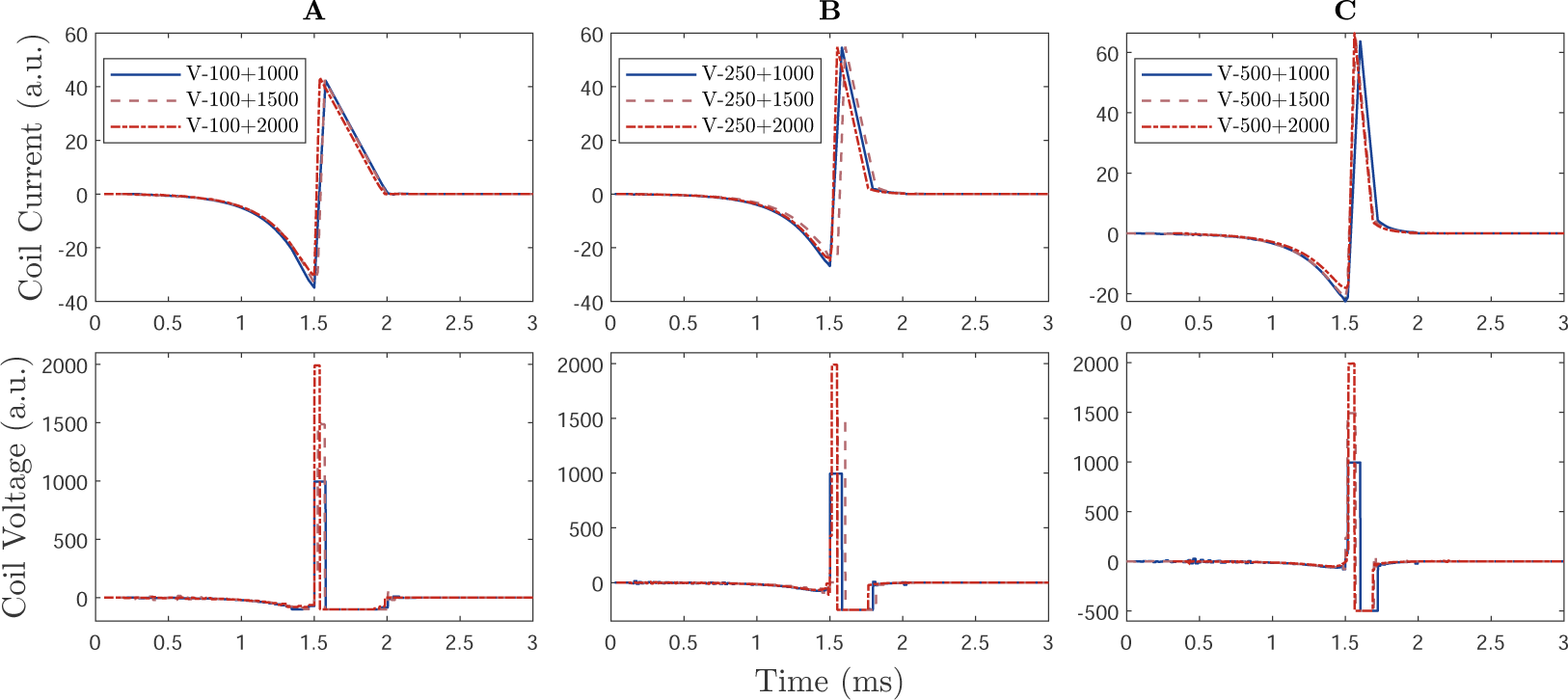
Voltage waveforms of the optimised current waveforms. Each column shows the waveforms with a given negative voltage limit and several positive voltage limits. a.u. refers to arbitrary units. (**A**) *V*_min_ = *−*100; (**B**) *V*_min_ = *−*250; and (**C**) *V*_min_ = *−*500. The upper row shows the coil current waveforms and the lower row shows the corresponding coil voltage waveforms. All of the coil voltage waveforms are nearly rectangular with different pulse duration and widths.

**Figure 3:**
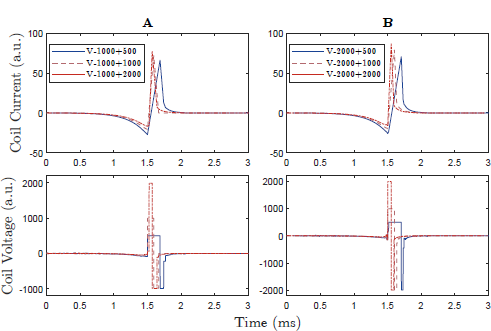
Voltage waveforms of the optimised current waveforms; a.u. refers to arbitrary units. Each column shows the waveforms with a given negative voltage limit and several positive voltage limits. (**A**) *V*_min_ = *−*1000; (**B**) *V*_min_ = *−*2000. The upper row shows the coil current waveforms and the lower row shows the corresponding coil voltage waveforms. All of the coil voltage waveforms are nearly rectangular with different pulse duration and widths.

### Current waveforms have dependencies on the pulse duration

The optimised results reveal several relationships between pulse shape and duration. Figures 4 (C) and (D) show that the peak current of the pulses at the threshold decrease for longer pulses while the depth of the current pre-phase increases. *Î*_max_, however, seems to approach a minimum, whereas the magnitude of *Î*_min_ increases without bounds. Accordingly, a deeper current pre-phase is needed to offset longer rising phases and pulse durations.

Figure 4 (E) shows that the pulse duration depends on the current ratio and the larger current ratio has a shorter pulse duration. In addition, because *Î*_max_ linearly depends on *Î*_min_ (shown in the inset plot), Figure 4 (E) also indicates that the rising phases of the current waveform are shifted upwards as the pulse duration decreases (also shown in Figures 2 and 3). With the purpose of minimising the energy cost, the phase duration should be shortened when the current amplitude is relatively high. Therefore, the larger current ratio correspondingly causes a shorter pulse duration, which, in turn, reduces the energy loss.

**Figure 4:**
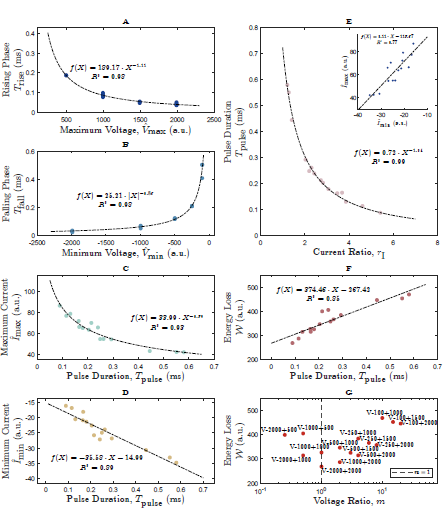
The relationship between the voltage amplitude and the corresponding pulse widths. a.u. refers to arbitrary units. (**A**) shows the positive voltage phases versus rising duration; (**B**) shows the negative voltage phases versus falling duration; (**C**) shows the pulse duration versus the maximum values of current waveforms; (**D**) shows the pulse duration versus the minimum values of current waveforms; (**E**) shows the current ratio *r*I versus pulse duration and the inset plot shows the linear relationship between *Î*_min_ and *Î*_max_; (**F**) shows the pulse duration versus the energy loss; (**G**) shows the voltage ratio versus energy loss and the vertical dashed line is *m* = 1 indicating symmetric field pulses. These coefficients of determination (*R*^2^ *≥* 0.77) suggest that these fitting curves accurately describe the trends.

Furthermore, the current waveform shape of the initial phase for asymmetric field pulses can be quantified. The initial phase follows a near-exponential course with a time constant of around 260 µs and a terminal value shifting the baseline for the subsequent phase dependent on the duration of the pulse duration (regression equations given in Figure 4). Thus, a quantitative equation for *Î*_min_ is

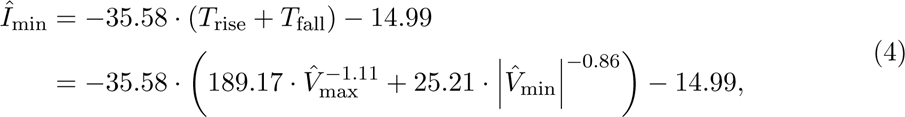

and initial phase duration *T*_init_ can be calculated by

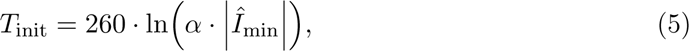

where *T*_init_ is in microseconds (µs) and *α* is an arbitrary and positive constant that controls the length of *T*_init_.

### The asymmetric field pulses can be more power-efficient than the symmetric field pulses

Figure 4 (F) illustrates that the energy loss is proportional to the pulse duration. These observations are consistent with Equation (1), which is the integral of the squared current over time. Moreover, higher voltage amplitudes lead to shorter pulse widths and short pulse duration and high pulse amplitudes have lower energy loss compared to longer ones. Considering the results given in Figure 4 (E), these results also suggest that the current waveforms with high current ratio *r*_I_ have low energy costs. Interestingly, Figure 4 (G) suggests that the optimised asymmetric field pulse could be more power-efficient than the symmetric ones. For example, the waveform of *V:–2000/+1000* has a lower energy loss than that of *V:–1000/+1000*. The waveforms with a positive voltage phase of 1000 consistently have higher energy loss compared to those with positive voltage phases of 2000 and 1500. Thus, these results further suggest that the limit of the positive voltage phase might determine the level of energy loss for a family of asymmetric field pulses, whereas the limit of the negative phase has a less pronounced effect.

### The asymmetric field pulses are more power-efficient than the monophasic pulses

Wang et al. (2023) recently optimised an original monophasic pulse used in MagPro X100 devices (MagVenture A/S, Farum, Denmark) and successfully reduced the energy loss by 12 % to 75 %. The authors formulated an optimisation problem that constrained the root-mean-square error (RMSE) of the optimised E-field pulse compared to the original monophasic pulse and the pulse duration. Although they forced the optimised pulses to stay close to the electric field pulse shape of a conventional monophasic device, some of the features such as the initial phase that shifts the current to an offset level from which the dominant pulse phase starts, also evolved there. To compare their results with ours, we selected the original pulse and their optimised pulses with RMSE values of 1 % and 4 % and pulse duration of 6 ms and scaled them such that they could successfully elicit an action potential in the neuron model. We searched the most similar current waveforms in our study to those of monophasic pulses by using cross-correlation measurements.

As shown in Figure 5, the original current waveform may correspond best to the current waveform of *V:–100/+2000*, the waveform with RMSE of 1 % to the current waveform of *V:–250/+1000*, and the waveform with RMSE of 4 % to the current waveform of *V:–500/+1500*. The asymmetric field pulses do not have high current amplitudes but the monophasic ones do. For example, the peak current of the original waveform is close to 70. In addition to the current waveforms, those previously optimised monophasic equivalents need higher coil voltage amplitudes compared to the optimised asymmetric field pulses with near-rectangular E-field pulses. As shown in Figure 5 (C), the voltage waveform with RMSE of 4 %, which allowed the optimisation of Wang et al. (2023) to deviate more from the conventional monophasic electric field waveform, has two sharp negative phases.

**Figure 5:**
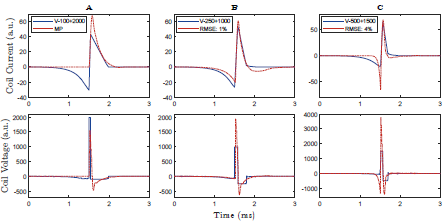
The optimised waveforms similar to the selected waveforms in Wang et al. (2023). a.u. means arbitrary unit. Three selected waveforms are (**A**) an original monophasic pulse (MP) recorded from a MagPro X100 device (MagVenture A/S, Farum, Denmark), (**B**) the optimised monophasic field pulses with root-mean-square error (RMSE) of 1 % and a pulse duration of 6 ms compared to the recorded one, (**C**) the optimised monophasic field pulses with RMSE of 4 % and a pulse duration of 6 ms compared to the recorded one.

As expected due to the fewer constraints and the priority of heating reduction in this paper, Figure 6 shows that the optimised asymmetric field pulses generally have a similar or lower energy loss compared to Wang’s optimised monophasic pulses, even though the monophasic equivalents share similar pulse duration with their corresponding asymmetric field pulses. Figure 4 (F) shows that many potential optimised waveforms in this study have lower energy loss than the monophasic pulses. These findings suggest that optimised near-rectangular asymmetric electric field pulses have an advantage over the more cosinusoidal monophasic pulses with a similar initial phase.

**Figure 6:**
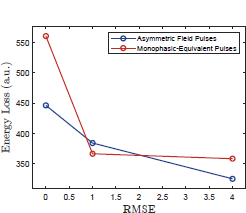
Comparison between the optimised monophasic-equivalent pulses reported in Wang et al. (2023) and the asymmetric field pulses in terms of energy loss. The x-axis represents the root-mean-square error (RMSE) constraint for the optimised monophasic pulses, and the y-axis the values of corresponding heating loss. Each optimised monophasic pulse (in red circle) has a similar asymmetric field pulse (in blue circle).

## 4 Discussion

In this study, we performed numerical optimisation of current waveforms for asymmetric field pulses, which are considered to offer a stronger activation selectivity than symmet-ric pulses, with the goal of minimising energy loss and coil heating while eliciting action potentials in neurons. Energy loss and coil heating were measured by the integral of the squared coil current. Our optimisation process was not limited to specific devices or pre-determined waveform parameterisation but rather a global search of the pulse shape space. We constrained the optimisation by imposing limitations on the maximum/minimum coil voltage and ensuring that the neuron model was able to generate an action potential. This approach allowed us to explore a wide range of potential waveform shapes and find solutions that are optimised for both low energy loss and effective neuron activation for various asymmetry ratios.

We used the terms asymmetric field pulse and symmetric field pulse to describe the triangle current waveforms that produce asymmetric and symmetric, near-rectangular electric field pulses, respectively. Traditional biphasic and monophasic pulses, on the other hand, induce underdamped or damped cosinusoidal electric field pulses. The optimal asymmetric current waveforms we identified consist of an initial slowly falling phase followed by rapidly rising and falling phases, trailed by a slow decay to zero. These features share similarities with the results for purely symmetric pulses reported by Goetz et al. (2013), part of which–specifically, the initial phase shifting the current baseline downwards–also evolved in Wang et al. (2023).

The current in the initial phase shifts the baseline so slowly that this phase hardly shows up in the electric field in most cases. The initial phase followed a near-exponential course with a time constant of around 260 µs for all optimised waveforms. The initial current phase, which has been reported previously from symmetric optimisation as well as from heat-reduced pulse-shape matching, appears as a major feature for most pulse shapes to reduce coil heating. The time constant of some 260 µs is above typical axonal time constants and the depth of the current pre-phase furthermore is so low that its electric field, i.e., the time derivative of the current, is only a fraction of that of the main phase (Barker 1991; D’Ostilio et al. 2016; Peterchev et al. 2013). Since this feature emerges many times and again in pulse-shape optimisation work, it appears that such a phase may be considered in pulses in general and could be added to pulse-shape designs provisionally without running computationally expensive numerical optimisation. Furthermore, our findings suggest that regardless of whether a pulse is asymmetric or symmetric, shorter pulses reduce coil heating and losses in the machine but require higher voltages to reach the threshold. This observation is in line with previous results for harmonic pulses (Barker et al. 1991; Cadwell 1991) and asymmetric field pulses (Peterchev et al. 2007, 2011).

The asymmetric field pulses would be useful in future TMS applications. Many studies have shown that some repetitive TMS protocols with monophasic or near-rectangular asymmetric pulses, such as quadri-pulse stimulation and 1 Hz repetitive TMS, can be more effective in producing sustained after-effects of the motor cortex than other protocols using biphasic pulses (Arai et al. 2007; Goetz et al. 2016; Hamada et al. 2007; Nakamura et al. 2016; Taylor and Loo 2007; Tiksnadi et al. 2020). Tiksnadi et al. (2020) suggested that the after-effect variability of the quadri-pulse stimulation with monophasic pulses was lower than that of the theta-burst stimulation with biphasic pulses. It is plausible to expect that the high-frequency or patterned protocols using asymmetric field pulses could potentially offer higher efficacy and lower variability, which would be beneficial for clinical applications because of their stronger selectivity of neuron populations.

Furthermore, recent studies have suggested that modifying the ratio between positive and negative electric field, i.e., the asymmetry level, and pulse widths of the asymmetric field pulse could further improve selectivity for specific neural populations while minimising energy loss compared to other pulses. For example, Sommer et al. (2018) suggested that the reverse phase of the asymmetric field pulse could prevent the activation of excitatory inputs to the corticospinal neurons and also recruit an additional population of excitatory inputs that are sensitive to the reverse phase. These results would be consistent with the idea that different current directions recruit distinct sets of excitatory synaptic inputs to corticospinal neurons (Di Lazzaro and Rothwell 2014). Moreover, TMS devices with adjustable waveforms have enabled the synthesis of a wide range of either rectangular field pulses with adjustable pulse duration, widths, and polarities (cTMS) or relatively freely designed pulses (modular pulse synthesizers) (Goetz et al. 2012a; Li et al. 2022; Peterchev et al. 2011; Zeng et al. 2022). Therefore, further optimisation of the asymmetric field pulse could enhance its selectivity and efficacy in targeting specific neural populations, which could have important implications for future TMS applications.

Although this study provided a theoretical framework for optimising the asymmetric field pulses, it is important to note that the optimal waveforms found are potentially only local minima and not necessarily global solutions. Furthermore, neurons and their ion-channel expression, as well as ion-channel accumulation along the axon are rather heterogeneous in the brain. Although nonlinear human neuron models may provide a relatively good approximation of average dynamical activation dynamics, no existing model can at present precisely simulate neural responses to time-varying electric fields. Therefore, fur-ther accuracy improvements in the optimisation process may require advancements in neural models or experiments. Despite these limitations, the stimulation results provide valuable insights into the underlying mechanisms and can serve as a starting point for experimental studies. The flexibility of the waveform representation and optimisation algorithm in this framework makes it suitable for alternative optimisation goals with appropriately formulated objectives in future waveform optimisation if quantitative models are available, such as stimulation selectivity, specific neural populations, or TMS sound reduction (Koponen et al. 2020; Peterchev et al. 2015b; Zhang et al. 2023).

## 5 Conclusion

Whereas previous optimisation aimed mostly at coil design, conventional symmetric electric field pulses, or almost perfect electric field equivalents of conventional monophasic pulses, thus introducing many constraints, we set up a minimally constrained optimisation routine to find pulses with minimum energy loss and coil heating within given practical voltage limits and various levels of asymmetry. Whereas at present clinical therapy uses practically exclusively symmetric biphasic pulses, which in the primary motor cortex show comparably low neuromodulatory strength, likely associated with their likewise lower activation selectivity, the asymmetric pulses studied here promise high efficacy. The reduced energy loss and coil heating of the asymmetric pulses are particularly attractive for future clinical use with high-rate repetitive stimulation or accelerated protocols. The lower energy loss and coil heating may further potentially allow reducing the cooling effort, which at present drive device cost.

For this purpose, we developed a hybrid global–local optimisation algorithm without constraining the pulse shape that produced a family of pulses, named asymmetric field pulse, with significantly lower energy loss compared to the traditional monophasic pulses. The asymmetric field pulses induce asymmetric and near-rectangular E-field by triangle current waveforms. We found quantitative levels for the initial current phase, which consistently had exponential shape with a time constant of around 260 µs so that it can likely be added to other pulse shapes as well to reduce coil heating. The optimised results also showed that the asymmetric field pulses could be more power efficient than symmetric ones.

## Author Contributions

SMG designed the research and initial method as well as code for minimally constrained optimisation of asymmetric field pulses. KM refined the method as well as the code and performed numerical optimisation. KM analysed the data and KM and SMG wrote the text.

## Notes

### Competing Interest Statement

The authors have declared no competing interest.

### Summary of Updates

This revision modified some inaccurate expressions, rearranged the plots, and fixed incorrect equations.

